# Self-organised attractor dynamics in the developing head direction circuit

**DOI:** 10.1101/221028

**Authors:** Joshua Bassett, Tom Wills, Francesca Cacucci

## Abstract

Head direction (HD) cells signal the orientation of an animal’s head relative to its environment. During post-natal development, HD cells are the earliest spatially modulated neurons in the hippocampal circuit to emerge. However, before eye-opening, HD cell responses in rat pups carry low directional information content and are directionally unstable. Using Bayesian decoding, we characterise this instability and identify its source: despite the directional signal being internally coherent, it consistently under-signals angular head velocity (AHV), incompletely shifting in proportion to head turns. We find evidence that geometric cues (corners) can be used to mitigate this under-signalling, and stabilise the directional signal even before eye-opening. Crucially, even when directional firing cannot be stabilised, ensembles of unstable HD cells show short-timescale (1-10 sec) temporal and spatial couplings consistent with an adult-like HD network, through which activity drifts unanchored to landmark cues. The existence of fixed spatial and temporal offsets across co-recorded cells and of an AHV-responsive signal, even before HD responses become spatially stable, suggests that the HD circuit is assembled through internal, self-organising processes, without reference to external landmarks. The HD network is widely modelled as a continuous attractor whose output is one coherent activity peak, updated during movement by angular head velocity (AHV) signals, and anchored by landmark cues. Our findings present strong evidence for this model, and demonstrate that the required network circuitry is in place and functional during development, independent of reference to landmark information.

## Introduction

Head direction (HD) cells are neurons that encode the direction of an animal’s bearing in the horizontal plane in rats [1] and in 3D coordinates in bats [2] in an allocentric (i.e. world-centred) reference frame. They are found in an extended cortical and subcortical network [3] and are the first among the hippocampal spatial networks to mature during postnatal development [4, 5]. The HD signal is commonly thought to support path integration [3] the ability of organisms to update their position by integrating angular and linear displacement [6, 7].

Simultaneously recorded HD cells in adult rodents maintain fixed offsets between their preferred firing directions under circumstances eliciting network re-orientation (e.g. changes to recording environment) [8, 9]. This coherence between HD cells is widely interpreted as evidence for a “ring attractor” network architecture, whereby continuous attractor dynamics allow a single peak of neural activity corresponding to HD within the ring, moved by inputs such as angular head velocity (AHV, signalling head turns) [10–12]. Evidence for the existence of attractor connectivity and AHV-like responses has recently been found in Drosophila [13, 14], while in the rat HD network, dynamics in sleep are also consistent with continuous attractor connectivity [15].

Currently, it is unknown when HD network architecture emerges during development and whether this process relies on learning (as suggested by models in which stable sensory input trains HD networks [16, 17]), or via experience-independent mechanisms.

Here, we set out to distinguish between these two possibilities and we find that, consistent with the latter scenario, attractor dynamics are present even before any stable HD tuning can be observed. Moreover, we identify one of the sources of instability in developing HD networks: systematic under-signalling of angular head velocity, resulting in catastrophic accrual of path integrative error. We also uncover how, during early post-natal life, geometric cues may be used to mitigate angular head velocity under-signalling and stabilise HD responses.

## Results

### Local cues can stabilise developing HD networks from post-natal day 13 onwards

Previous studies have shown that HD cells can be recorded before eye opening in the rat, but pre-visual HD cells are few in number, carry low levels of directional information and exhibit spatial instability, compared to those of adults or post-eye opening pups [18, 19]. Tan et al (2015) recorded HD cells from rat pups moving freely within a 62 × 62cm open field environment (“standard box”) typical of that used with adult animals [4, 18]. Here, we also tested responses introducing animals in a second, smaller environment (“small box,” 20 × 20cm), limiting the exploratory range. We recorded in the anterior thalamus (1276 cells from 21 animals, aged Post-natal day (P) 12-21) while rat pups alternately explored the two environments (Figure 1A). The proportion, spatial tuning and stability of HD cells were higher in the small vs the standard box (Figure 1B), most markedly at the pre-visual ages P13-14 (2-sample Z-test for proportions, Z>7.5, p<0.0001 for both P13-14; ANOVA RV, Age*Env F_4,1025=15_, p<0.001, Simple Main Effects (SME) (ENV) p<0.001 at P13-14; ANOVA stability, Age*Env F_4,1019_=45, p<0.001, SME_(ENV)_ p<0.001 at P13-14), indicating that sensory modalities other than vision are capable of anchoring HD representations before eye opening.

**Figure 1.**
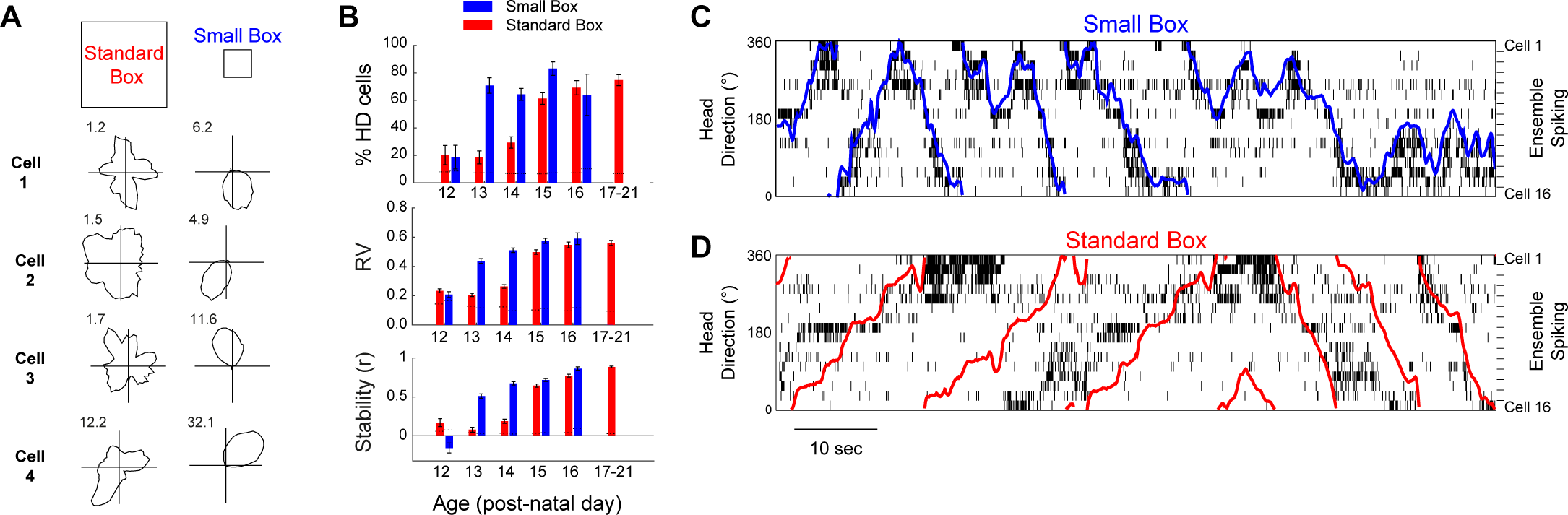
HD cell signalling is stabilised when rats explore a small (20cm side) environment, from P13 onwards. (**A**) Example firing rate polar plots for 4 HD cells recorded at P13 in the standard (left) and small (right) box. Numbers top left indicate peak firing rate (Hz). (**B**) HD cells are more numerous (top), have a higher spatial tuning (Rayleigh vector [RV], middle) and intra-trial stability (bottom) when recorded in small vs standard box in animals older than P13. (**C, D**) HD cell network internal organisation is preserved in the standard box, even when directional signalling is unstable. Coloured traces indicate the rat’s actual head direction, overlaid upon spike raster plots for all simultaneously recorded HD cells, in the small (C) and standard (D) boxes. For both (C) and (D) HD cells are ordered vertically by their preferred firing direction in the small box. The sequences of HD cell activation for each head turn direction are similar in the small and standard boxes, but the direction signalled by HD cell firing consistently undershoots actual rotation, in the standard box.

At P13-P14, the internal organisation of the HD network appears to be preserved even when HD responses are unanchored to the laboratory frame of reference (the standard box): for each head turn direction, similar spiking sequences can be observed across co-recorded HD cells in both small and standard boxes. Network activity transitions smoothly through directional space, whilst often undershooting actual head movements (Figure 1C).

### HD cells display adult-like spatial and temporal coupling, even when drifting

In order to confirm whether the internal organisation of the HD network is preserved at these ages, even when its responses are unanchored from the external environment, we examined the short-term temporal and spatial couplings between pairs of co-recorded neurons. We computed temporal cross-correlograms (Figure 2A and 2B) and time-windowed spatial cross-correlograms (Figure 2D and 2E) for all pairs of co-recorded cells which displayed HD tuning in the small box (see Methods). In order to eliminate residual HD stability in the standard box as a confounding factor we excluded any cells which displayed significant HD tuning in the standard box. Despite this, both the temporal and spatial relationships between pairs of co-recorded HD cells are preserved across both the small and standard environments (significantly correlated temporal and spatial offsets between small and standard box, see Figure 2C and 2F). This demonstrates that even when HD cells are unstable in the open field (on P13-14), the internal network structure is unchanged, compared to when HD signalling is stable.

**Figure 2.**
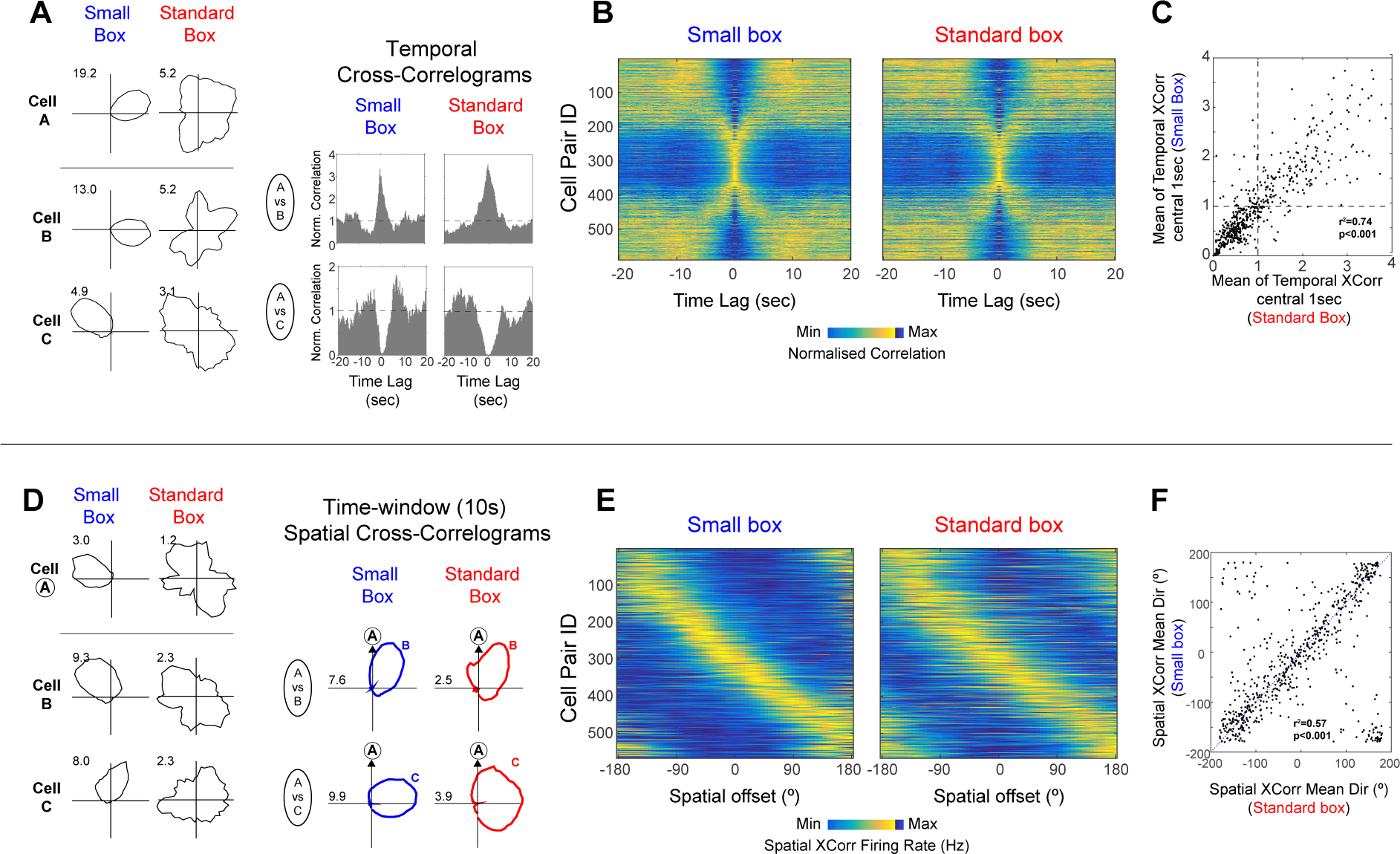
Short-time scale temporal and spatial couplings between HD cells are preserved even when directional signalling is unstable. (**A**) Example polar plots (left) and temporal cross-correlograms (TXCs, right) for 3 HD cells recorded in small and standard box. (**B**) TXCs of all co-recorded HD cell pairs in the small (left) and standard (right) box, normalised between their minimum (dark blue) and maximum (yellow) values. Each row in the image shows one TXC, rows are sorted on the basis of preferred firing direction difference in the small box. (**C**) Correlation between mean values of the central 1 sec of TXCs in the small vs standard boxes. (**D**) Example polar plots (left) and time windowed (10 sec) spatial cross-correlograms (SXC, right) for 3 HD cells. (**E**) SXCs of all co-recorded HD cell pairs in the small (left) and standard (right) boxes. HD pairs sorted as in (D). (**F**) Circular-circular correlation between the mean directions of SXCs in the small vs standard boxes.

### Attractor connectivity precedes HD landmark stabilisation

Introducing the youngest group of animals (P12) in the small box did not result in an improvement in HD cell stability/tuning (Z-test for % HDC, Z=0.71, p=0.45; SME_(ENV)_ for RV, p=0.99; for stability, small box significantly lower, SME_(ENV)_ p=0.002; Figure 1B). Nevertheless, many co-recorded cell pairs showed temporal and spatial coupling indicative of an attractor neural structure, even at this age (Figure 3A, 3B and 3C). To investigate whether the degree of temporal and/or spatial coupling at P12 was significantly higher than chance, we measured the proportions of P12 cross-correlogram scores lying beyond 95% confidence limits for HD-HD coupling (defined as the 5^th^ and 95^th^ percentiles of the scores from all known non-HD cell pairs in older rats; figure 3d; 95^th^ percentile only for spatial cross-correlogram RV). The proportion of P12 cell pairs with coupling scores beyond these confidence limits significantly exceeds 5% (one sample Z-test; temporal lower tail Z=19, p<0.001, temporal upper tail Z=14, p<0.001, spatial upper tail Z=33, p<0.001), demonstrating that HD neurons display fixed temporal and spatial offsets even at P12, an age at which none of the tested experimental manipulations results in environmentally stable HD responses. The HD network thus displays a key signature of continuous attractor structure before HD responses can be stabilised by local landmarks.

**Figure 3.**
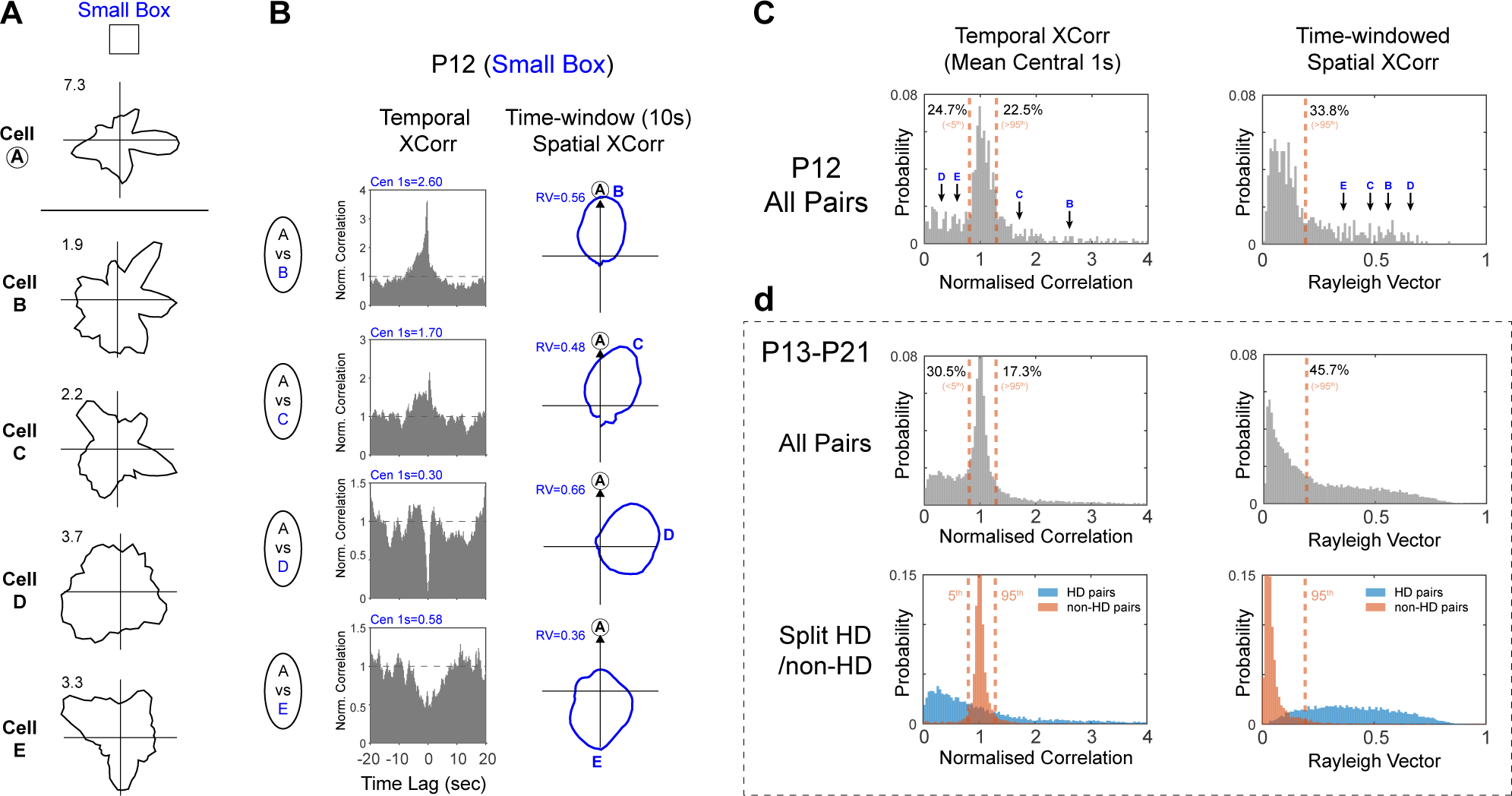
HD attractor network organisation is present at P12, even before HD cells can be anchored to landmarks. (**A**) Polar plots for five example cells recorded in small box at P12, showing no HD tuning over a 10 min session. (**B**) Temporal cross-correlograms (TXCs; left) and spatial cross-correlograms (SXCs; right) between Cell A and Cells B-E. Blue text top-left shows mean of central 1 sec of TXC and RV length of SXC, respectively, for each example. (**C**) Probability distributions of mean of central 1 sec of the TXC (left) and RV length of the SXC (right) scores for all P12 co-recorded cell pairs (N=452). Black arrows with blue letters indicate scores from examples shown in (A, B). Orange dashed lines show the values of the 5^th^ and 95^th^ percentiles of scores of known non-HD cells in older animals (see (D); only 95^th^ percentile is shown for SXC). Black text refers to percentages of P12 scores above or below these percentiles. (**D**) Top row: as for (C) but for all co-recorded cell pairs P13-P21. Bottom row: same data as top row but distributions of HD-HD pairs (light blue) and nonHD-nonHD pairs (orange) plotted separately. Orange dashed lines shown here and in (C; D top row) are derived from the 5^th^ and 95^th^ percentile of the orange (nonHD-nonHD) distributions.

### Drift of the HD representations occurs due to systematic under-signalling of angular head velocity

In order to further characterise the temporal dynamics of drifting HD networks, we applied a cross-trial Bayesian decoding approach, using firing rate maps of stable HD cells in the small box to decode signalled direction (in small box coordinates, see Methods) as the rat moved in the standard box, at P13-14. Although, as expected, decoded headings diverged from actual headings, decoding produced a coherent estimate of direction exhibiting continuous, smooth transitions during a trial, consistent with attractor dynamics (Figure 4A, top; Figure S1). The coherence and smoothness of decoded trajectories were not significantly different whether decoding was performed on standard or small box data, further demonstrating the maturity of the internal network dynamics, despite ongoing drift (figure S1E-F). We then obtained the decoded angular head velocity by calculating the first derivative of the decoded head direction (AHV, Figure 4A, bottom; see Methods). We found that, when HD responses are drifting, although AHV is linearly correlated with actual head velocity, it is under-signalled by drifting HD networks in the standard box (Figure 4B regression β significantly lower for standard than small box, t_(6916)_ = 16.54, p<0.0001). Thus, the most prominent source of error in young HD networks is a systematic under-signalling of AHV, which cannot be compensated for by alternative sensory cues in the standard box. No significant differences between angular velocity profiles in small and large box can be detected at any age (see Figure S2), discounting this as a potential factor accounting for the differential AHV error accumulation across these conditions.

**Figure 4.**
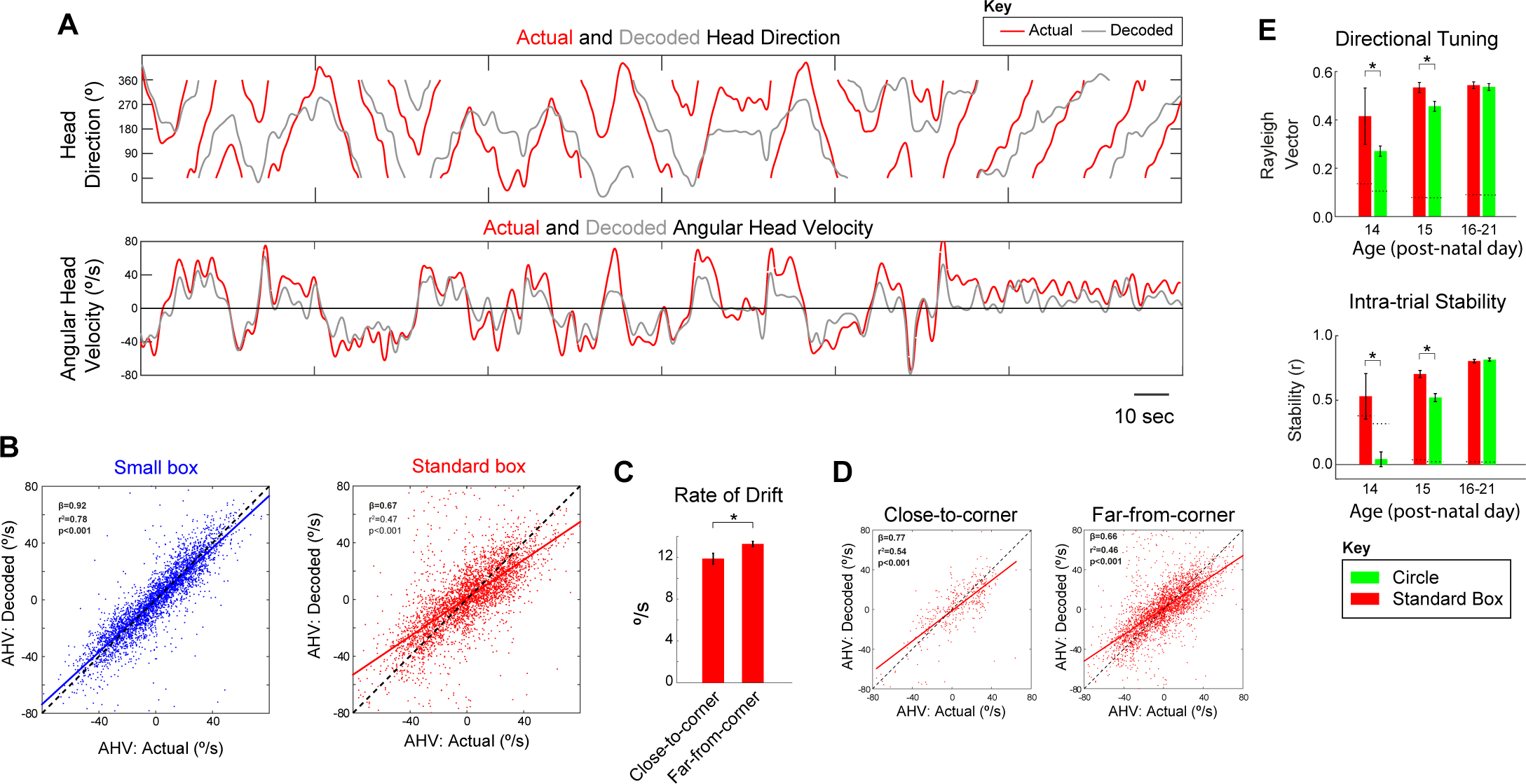
HD drift in young rodents is caused by angular head velocity (AHV) under-signalling, and is reduced by proximity to corners. (**A**) Example of actual (red) and decoded (grey) head direction (top) and angular head velocity (bottom) values displayed by a P14 rat during 5 minute exploration in the standard box. (**B**) Correlation between actual (x-axis) and decoded (y-axis) angular head velocity scores in the small (left) and standard (right) boxes across all decoded ensembles (N=6). Slope of relationship between actual and decoded AHV is significantly smaller in the standard vs small box. (**C**) Mean (±SEM) rate of drift (rate of divergence between actual and decoded head direction) when rats were close or far from the corners of the standard box. (**D**) AHV under-signalling is reduced when rats are close to corners. Correlations between actual and decoded AHV scores in the standard box, split by proximity to corners (left, close; right, far). (**E**) Directional tuning and intra-trial stability of HD cells are reduced in a circular environment, as compared to the standard (square) box, on P14–15. Bar charts show the mean (±SEM) Rayleigh vector (top) or intra-trial stability (bottom) of HD cells recorded in standard and circular environments.

**Supplemental Figure 1.**
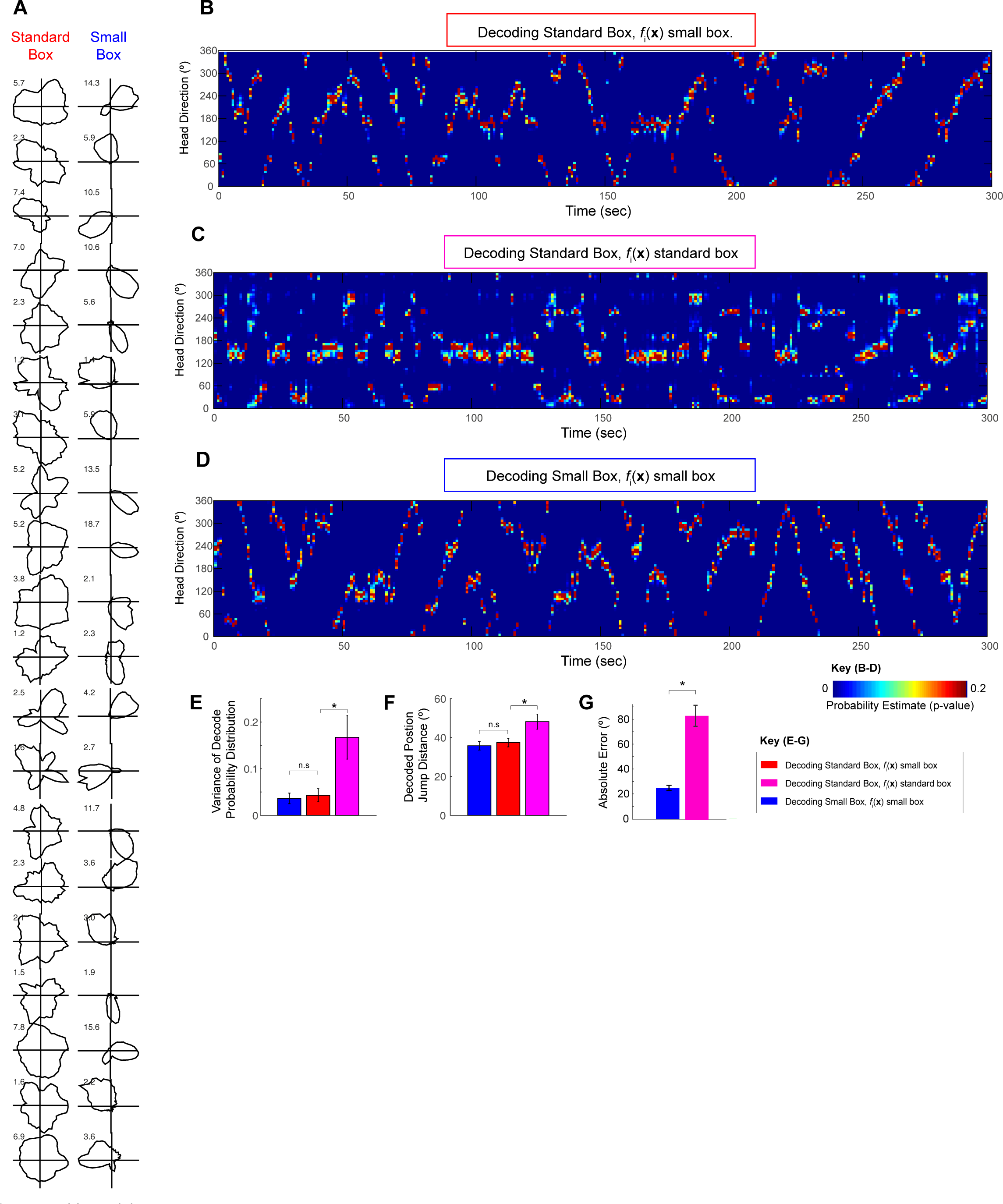
Validation of cross-trial (small/standard box) Bayesian decoding approach at P13-P14. (**A**) Polar plots for all HD cells used for decoding in the example of main figure 2a, recorded in standard box and small box (P14). HD cells show stable directional signalling in the small box (right), but not in the standard box (left). (**B**) Full set of decoded probability estimates for example shown in main figure 2a. False colour image showing probability estimates of the rat facing in each direction (y-axis, bin size 6° width), computed from HD spiking in each decoding window (x-axis, window size = 1 sec). The function of average firing in a given direction for each cell (f_i_(x) where i represents the cell and x the direction) was based on polar plots derived from the small box trial. A coherent cluster of high probability is apparent in each decoding window and the estimated direction (maximum probability) moves smoothly between time bins, consistent with the presence of continuous attractor dynamics in the HD cell network, even though the HD network is not anchored to laboratory reference frame (see polar plots in (A) left column). (**C**) Full set of probability estimates obtained by decoding the same data as (A) and main figure 2a, in the case where f_i_(x) is based on the polar plots derived from the standard box trial. Consistent with the reduced spatial tuning and stability of the polar plots in the standard box, distributions of probability estimates within each decoding window display a high variance, and discontinuous jumps between temporally contiguous decoding windows are apparent. (**D**) 5 minute example of full probability estimates obtained decoding the small box trial (polar plots in (A), rightmost column). f_i_(x) was based on data derived from the same small box trial data (interleaved 60-sec time epochs were used for f_i_(x) construction and decoding, respectively, see methods for details). Comparison with (B) shows that the circular coherence of the probability estimates and the smoothness of the decoded trajectory are qualitatively similar whether decoding the small box, or the standard box using small box f_i_(x). (**E**) For all decoded ensembles (N=6), there is no significant difference in the circular coherence of probability within each window (measured by weighted circular variance), when comparing small box decoding and standard box decoding using small box f_i_(x). By contrast, standard box decoding using standard box f_i_(x) leads to significantly greater variance (ANOVA; F_2,15_=6.51, p=0.009; Post-hoc HSD; small vs standard f_i_(x) small, p=0.98, standard f_i_(x) small vs f_i_(x) standard, p=0.021). (**F**) For all decoded ensembles, there is no significant difference in the smoothness of decoded trajectories (measured by the mean angular jump between consecutive decoded directions), when comparing small box decoding and standard box decoding using small box f_i_(x). By contrast, standard box decoding using standard box f_i_(x) leads to significantly greater average jumps (ANOVA; F_2,15_=5.6, p=0.015; Post-hoc HSD; small vs standard f_i_(x) small, p=0.91, standard f_i_(x) small vs f_i_(x) standard, p=0.043). (**G**) Spatial accuracy of decoding, measured by absolute distance between actual and decoded head direction. Decoding the small box leads to absolute errors consistent with previously published studies^14^. By contrast, absolute error in the standard box is larger when using small box f_i_(x), indicating that the HD cells are not stably anchored to the testing environment reference frame (T-Test, t_5_=6.1, p=0.002).

**Supplemental Figure 2.**
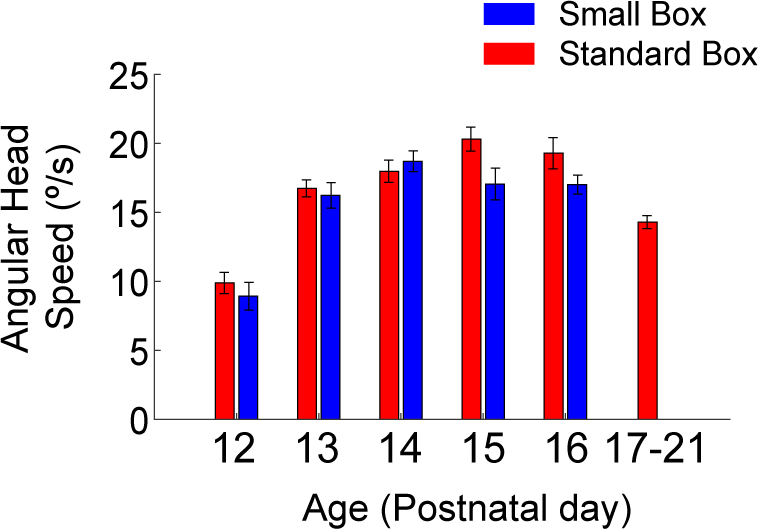
Head angular speed (defined as the unsigned angular head velocity) does not differ significantly between the small and standard boxes, between P12 and P16. (**A**) Overall mean angular speed (±SEM) in standard and small box at each age. No significant differences between angular speed in small and large box at any age can be detected (2-way ANOVA Age*Env; Env F_1,294_=2.86, p=0.09, Age F_1,294_=21.8, p<0.0001, Env*Age F_4,294_=1.47, p=0.21).

### Geometric cues can mitigate path integration error

Introducing rat pups in the small box results in stabilisation of HD responses (Figure 1A and 1B) and reduction of angular head velocity under-signalling (Figure 4B, left). Following previous work showing that boundaries stabilise place cell firing during post-natal development [20], and grid cell signalling in adult mice [21], we tested whether HD cells were more stable when animals were close to an environmental boundary. Unexpectedly, the rate of drift (rate of divergence between actual and expected heading) was significantly greater close to walls (Mean Rate of Drift: walls=13.3±0.52°/s, centre=12.2±0.70°/s; t-test: t_(3476)_=2.5, p=0.007). This is likely caused by an increase in actual AHV when animals are close to walls (Mean AHV: walls=21.6±0.6°/s, centre=20.2±0.8°/s; t-test: t_(3476)_=2.70, p=0.012). However, further analyses showed that, interestingly, both the rate of drift of HD responses and AHV under-signalling were significantly reduced when animals were close to a corner (Mean Rate of Drift: t-test: t_(3476)_=1.98, p=0.048; see Figure 4C; AHV under-signalling: t-test for difference of β: t_(3476)_=2.47, p=0.013, see Figure 4D). This effect cannot be accounted for by differences in actual AHV close to and far from corners (Mean actual AHV: corners=20.1±0.7, far-from-corners=20.9±0.2; t-test: t_(3478)_=0.97, p=0.33). Corners may therefore be a geometric feature used to offset path integration error in developing animals.

To further investigate the stabilising influence of corners on the developing HD network, we introduced rat pups (P14-15) in a circular environment of similar dimensions to the standard square environment (matched perimeter). When recorded in the circular environment, HD cells showed reduced directional tuning and intra-trial stability (Figure 4E; Directional tuning [RV]: ANOVA Age*Env F_(2,348)_=8.5, p<0.001, SME_(ENV)_: P14, p=0.023, P15, p<0.001. Stability: ANOVA Age*Env F_(2,343)_=18.9, p<0.001, SME_(ENV)_: P14, p<0.001, P15, p<0.001). These findings further support the interpretation that corners may be used to stabilise HD cells in early development.

### Discussion

The HD circuit has been extensively modelled as possessing a 1-dimensional ring architecture, with connectivity between units arranged such that nodes with similar heading preferences are mutually excited, whereas nodes with widely diverging preferred head directions inhibit each other [10–12]. This topology gives rise to continuous attractor dynamics, whereby the network always settles on one solution, resulting in a single, localised peak of activity, involving neighbouring nodes. This solution represents current head bearing, and is thought to be updated in response to head movement through the activity of angular velocity responsive cells, which are asymmetrically connected to HD neurons. Angular velocity correlated responses have been described in the rat midbrain [22, 23], and in the central complex of *Drosophila* [14, 24]. Thus, in this model, the HD network performs a velocity-to-position integration, a computation sufficient to support the angular component of path integration behaviours. Indeed, lesions of the HD system significantly impair performance on spatial tasks that depend on angular path integration [25, 26].

Path integration of any kind is prone to overwhelming accumulation of error without reference to landmark cues that allow resetting of the integration process, effectively discharging the error and anchoring the HD representation to the external environment [10–12, 27]. If the HD system is both a ring attractor and a path integrator, then the developing HD network requires not only the intrinsic ring topology, but a self-motion input (e.g., angular head velocity) and landmark cues corresponding to stable features of the environment.

To date no experimental evidence addresses how or in what order these three requirements are met during brain development, but theoretical models of the emergence of the HD system suggest the presence of a distal landmark cue which acts as a supervisory input, simultaneously anchoring the network to the external world and instructing the correct wiring across nodes [16, 17].

Here, we show that during early post-natal development (P12-14), days before eye-opening, when rat pups explore open field environments, the internal dynamics of the network are exactly as described in a mature continuous attractor, even while its spatial relationship to the environment is adrift. Using Bayesian decoding we have revealed the temporal dynamics of this drift and, in doing so, provided further strong evidence that the mammalian HD network shows a key signature of attractor dynamics: smooth, continuous transitions across network states even when the system is not responding to landmark cues [15]. Furthermore, we have shown for the first time that these network dynamics are present less than two weeks after birth. Our results provide new constraints on models of continuous attractor network development, as they demonstrate that the rigid coupling of spatial relationships across directional nodes is not extracted from the structure of sensory input through a learning process [16, 17], but seem to emerge, at least in part, via experience-independent processes [28]. Taken together, our results provide strong experimental support for the existence of ring attractor architecture of HD networks in the rat, and demonstrate that this likely arises through internal, self-organised processes during development. This finding may also generalise to other putative attractor networks such as grid cells [29–32], which display strong coupling as soon as they can be detected in young animals [33].

Our analysis of the temporal dynamics of the spatial drift observed in pre-eye-opening pups’ HD cells has allowed identification of its underlying source: the HD attractor in young animals shows a linear response to angular head velocity, but with reduced gain, indicating systematic under-signalling. This constitutes direct experimental evidence that the attractor’s activity peak is shifted around the ring at displacement values proportional to the magnitude of angular head velocity input [34], thereby integrating velocity to position as predicted by current models of the HD network. In the open field, the under-signalling of velocity, combined with the inability to correct these errors using landmarks amounts to an open-loop state, resulting in the observed drift of directional firing with respect to the external environment. It remains undetermined whether the ultimate cause for this under-signalling can be imputed either to reduced vestibular input [27], to its incomplete integration by the HD network, or a combination of the two. Further work mapping the development of angular head velocity responses across the sub-cortical HD circuit would help to dissociate between these possibilities.

We have identified that corners are important landmark cues that may be used by the developing HD network to discharge path integrative error. Our results suggest that geometrical features, sensed through olfactory/tactile modalities, can be used to stabilise HD responses during early post-natal development, at a time when access to distal cues is still limited. These results provide new mechanistic insights as to the recognised importance of geometric features in the development of spatial cognition in children [35, 36].

## ACKNOWLEDGEMENTS

We acknowledge funding from the ERC (‘DEVSPACE’ grant to FC and JPB position), the HBP, the Medical Research Council, UK, and the Royal Society (URF fellowship to TW). We would like to thank the following for helpful discussion of previous versions of this manuscript: Caswell Barry, Andrej Bicanski, Neil Burgess, Ila Fiete, Andreas Herz, Kate Jeffery, Colin Lever and Hector Page.

## Author contributions

JPB, FC and TW conceived the experiments, and JBP and FC conducted them. TW analysed the data with contributions from FC and JPB; FC, TW and JPB wrote the manuscript.

## Figure legends

**Figure 1**. HD cell signalling is stabilised when rats explore a small (20cm side) environment, from P13 onwards. (**A**) Example firing rate polar plots for 4 HD cells recorded at P13 in the standard (left) and small (right) box. Numbers top left indicate peak firing rate (Hz). (**B**) HD cells are more numerous (top), have a higher spatial tuning (Rayleigh vector [RV], middle) and intra-trial stability (bottom) when recorded in small vs standard box in animals older than P13. (**C, D**) HD cell network internal organisation is preserved in the standard box, even when directional signalling is unstable. Coloured traces indicate the rat’s actual head direction, overlaid upon spike raster plots for all simultaneously recorded HD cells, in the small (C) and standard (D) boxes. For both (C) and (D) HD cells are ordered vertically by their preferred firing direction in the small box. The sequences of HD cell activation for each head turn direction are similar in the small and standard boxes, but the direction signalled by HD cell firing consistently undershoots actual rotation, in the standard box.

**Figure 2**. Short-time scale temporal and spatial couplings between HD cells are preserved even when directional signalling is unstable. (**A**) Example polar plots (left) and temporal cross-correlograms (TXCs, right) for 3 HD cells recorded in small and standard box. (**B**) TXCs of all co-recorded HD cell pairs in the small (left) and standard (right) box, normalised between their minimum (dark blue) and maximum (yellow) values. Each row in the image shows one TXC, rows are sorted on the basis of preferred firing direction difference in the small box. (**C**) Correlation between mean values of the central 1 sec of TXCs in the small vs standard boxes. (**D**) Example polar plots (left) and time windowed (10 sec) spatial cross-correlograms (SXC, right) for 3 HD cells. (**E**) SXCs of all co-recorded HD cell pairs in the small (left) and standard (right) boxes. HD pairs sorted as in (D). (**F**) Circular-circular correlation between the mean directions of SXCs in the small vs standard boxes.

**Figure 3**. HD attractor network organisation is present at P12, even before HD cells can be anchored to landmarks. (**A**) Polar plots for five example cells recorded in small box at P12, showing no HD tuning over a 10 min session. (**B**) Temporal cross-correlograms (TXCs; left) and spatial cross-correlograms (SXCs; right) between Cell A and Cells B-E. Blue text top-left shows mean of central 1 sec of TXC and RV length of SXC, respectively, for each example. (**C**) Probability distributions of mean of central 1 sec of the TXC (left) and RV length of the SXC (right) scores for all P12 co-recorded cell pairs (N=452). Black arrows with blue letters indicate scores from examples shown in (A, B). Orange dashed lines show the values of the 5^th^ and 95^th^ percentiles of scores of known non-HD cells in older animals (see (D); only 95^th^ percentile is shown for SXC). Black text refers to percentages of P12 scores above or below these percentiles. (**D**) Top row: as for (C) but for all co-recorded cell pairs P13-P21. Bottom row: same data as top row but distributions of HD-HD pairs (light blue) and nonHD-nonHD pairs (orange) plotted separately. Orange dashed lines shown here and in (C; D top row) are derived from the 5^th^ and 95^th^ percentile of the orange (nonHD-nonHD) distributions.

**Figure 4**. HD drift in young rodents is caused by angular head velocity (AHV) under-signalling, and is reduced by proximity to corners. (**A**) Example of actual (red) and decoded (grey) head direction (top) and angular head velocity (bottom) values displayed by a P14 rat during 5 minute exploration in the standard box. (**B**) Correlation between actual (x-axis) and decoded (y-axis) angular head velocity scores in the small (left) and standard (right) boxes across all decoded ensembles (N=6). Slope of relationship between actual and decoded AHV is significantly smaller in the standard vs small box. (**C**) Mean (±SEM) rate of drift (rate of divergence between actual and decoded head direction) when rats were close or far from the corners of the standard box. (**D**) AHV under-signalling is reduced when rats are close to corners. Correlations between actual and decoded AHV scores in the standard box, split by proximity to corners (left, close; right, far). (**E**) Directional tuning and intra-trial stability of HD cells are reduced in a circular environment, as compared to the standard (square) box, on P14-15. Bar charts show the mean (±SEM) Rayleigh vector (top) or intra-trial stability (bottom) of HD cells recorded in standard and circular environments.

## Methods

### Subjects

Subjects were 21 male Lister Hooded rats (RGD_2312466) aged P12-P21 weighing 18-29g at the time of surgery. Litters were bred on site and implanted subjects remained with their mothers and litter-mates throughout the experimental period. Litters were housed in 42×32×21 cm cages furnished with nesting material and environmental enrichment objects, and maintained on a 12:12 hour light:dark schedule with lights off at 13:00. Litters were culled to 8 pups at P4 in order to minimise inter-litter variability. Implanted pups were separated from the litter for between 20 to 120 minutes per day for electrophysiological recordings.

### Surgery and electrodes

Rats were anaesthetised using 1-3% isoflurane and buprenorphine via subcutaneous injection at 0.15mg/kg of body weight. Rats were implanted with 8 tetrodes consisting of HM-L coated 90% platinum/10% iridium 17μm wire (California Fine Wire, Grover City, CA). The implanted apparatus weighed 1 gram. Tetrode bundles were implanted into the ADN using the following stereotaxic coordinates: 1.7 mm posterior to bregma, 1.2mm lateral from the midline at bregma, and 4.2mm ventral from the skull surface at bregma. Following surgery, rats were placed on a heating pad until they could move spontaneously and then were returned to the home cage. After experiments were completed, tetrode position was confirmed by transcardially perfusing the rat (4% formaldehyde in PBS) whilst the tetrodes remained in their final position, followed by brain sectioning at 30μm, and Nissl-staining of the resulting sections.

### Single-unit recording

Following surgery, rats were allowed 24 hours recovery. Tetrode bundles were then advanced ventrally in increments of 62.5-250 μm/day. Experimental recording sessions began when any single unit neural activity could be identified. Single unit data was acquired using the DACQ system (Axona Ltd, St. Albans, UK). Position and directional heading was recorded using a 2-point tracking system consisting of 2 LEDs spaced 7 cm apart and attached to the headstage amplifier in a fixed orientation relative to the animals’ head. Isolation of single units from tetrode-recorded data was performed manually on the basis of peak-to-trough amplitude or principal components, using the TINT software package (Axona Ltd., St Albans, UK) with the aid of KlustaKwik1 automated clustering (1).

### Behavioural Testing

Single-unit recording trials took place in one of two square recording arenas. The ‘standard box’ had a side length of 62.5cm and was 50cm high, painted light grey, and placed on a black platform. The box was placed in the open laboratory, and distal visual cues were available in the form of the fittings and contents of the laboratory. The floor of the arena was not cleaned. There were no further polarising cues placed within the recording arena. In order to ascertain whether cells which did not display a stable HD correlate in the standard box could be anchored to the laboratory frame in any other environment, we recorded the activity of the cells in a smaller square box (‘small box’: 20cm side length, 42cm high). The small box was placed on the same platform as the standard box and centred in the same location. In the majority of small box trials, the small box contained two polarising cues, a 3D piece of wood (2cm × 4cm × 42cm), placed in the NW corner, and a sheet of polystyrene covering the E wall. Rats were subject to between 1 and 4 standard box trials (median 2), and 1 to 5 small box recording trials (median 2) per session (maximum total trials 7, median 3). As HD cells in the standard box were already mature by P15-16 (see also [2–4]), the oldest rats (P17-21) were not tested in the small box. Rat pups were kept in a separate holding box (40 × 40 × 25cm) furnished with bedding and a heating pad in between recording trials. A sub-set of rats (N=14, contributing 351 HD cells) were also tested in a circular environment, from P14 onwards. The circular environment was 79cm diameter, and (similarly to the standard box) was wooden, painted light grey, and placed on a black platform. A large plain white cue card (75cm × 1m), illuminated by a 40W lamp was placed outside the environment, approximately 75cm away from its edge. No polarising cues were placed inside the environment.

### Construction of firing rate maps

To minimise artefactual correlates due to under-sampling of position, data was only included if the angular path length for the session exceeded the equivalent of 25 full head turns (values derived from the 5^th^ percentile of the whole dataset). Positional (directional) data was sorted into 6° bins in the yaw plane. Following this, total dwell time, d, and spike count, s, for the whole trial was calculated for each directional bin. The binned position dwell time and spike count maps were then smoothed using a 30° boxcar filter, and the rate for each directional bin is defined as s/d.

### Classification of single-units as HD cells

To minimise artefactual correlates due to under-sampling, only cells which fired at least 100 spikes in a recording session were included in further analysis. Single units were classified as HD cells if the mean resultant vector length (Rayleigh vector; RV) of the directional firing rate map exceeded a threshold defined as the 95th percentile of a population of RV scores derived from age- and brain area-matched spatially shuffled data (2). Briefly, shuffled data was generated by shifting spike trains relative to position by a random amount between 20 seconds and trial duration minus 20 seconds, leaving the temporal structure of the spike train and the positional data otherwise unchanged. The shuffled data was then used to construct directional rate maps, as described above. This process was repeated a sufficient number of times for there to be 100,000 shuffled RV values for every 1-day age group, for each brain area. Single units with an RV ≥ 95th percentile of this shuffled population were defined as HD cells.

### Quantitative analysis of directional signalling

‘*% HD cells*’ was defined as the number of neurons classified as HD cells divided by number of total cells recorded in an environment (standard or small box). The difference between proportions of HD cells in the standard and small boxes in each age bin was tested using a 2-sample Z-test for proportions. ‘*Intratrial stability*’ was defined as the correlation between spatially corresponding bins from the first and second half of a single trial, using only those bins in which firing rate > 0Hz in at least one half of the trial. The effects of age and environment on intra-trial stability and the RV of HD cells were tested initially by a 2-way ANOVA, followed by tests of simple main effects to assess differences between environments at particular ages.

### Temporal and Spatial Relationships between Cells

Temporal cross-correlograms were defined as the cross-correlograms (constructed using the matlab ‘xcorr’ function) between time-binned histograms of the spike times of two neurons (bin width = 200ms). In order to standardise the height of the central peak (or depth of the central trough) and allow comparisons across all cell pairs, raw temporal cross-correlograms were normalised by the mean of a population of cross-correlograms (100 per pair) constructed with spike-shifted data. The 100 spike-shifted correlograms were constructed by shifting the spikes of one neuron along a set of linearly spaced intervals from 20sec to [trial duration - 20sec]. The mean normalised spike count of the central 1 second of the cross-correlogram was then used as a measure of the degree of offset (i.e. coupled or anti-coupled firing) between neurons. The degree of similarity between central 1-sec spike counts in the large and small boxes was compared using linear regression. Only cell pairs in which both contributing cells were classified as HD cells in the small box, and neither were classified as HD cells in the standard box, were included in this analysis.

Time-windowed spatial cross-correlograms between pairs of neurons were constructed analogously to those in (5, 6) but with spatial firing being assessed with respect to head direction, rather than to 2-dimensional space. The spike times of one neuron (the ‘reference’ neuron) were used to define a series of 10 sec time windows, each starting at the time of a reference neuron spike (head directions of ‘reference neuron’ were assigned a zero degrees value - windows were reduced in duration if a reference spike occurred within 10 secs of the beginning or end of the trial). Within each window, a histogram was created of the head directions associated with the spiking of the second neuron (the ‘test’ neuron; histogram bin width = 6°), with directions being defined relative to the window centre. The histograms of all windows were then summed, and smoothed (30° boxcar filter), producing an overall map of the directions in which the test neuron fired relative to the reference neuron, within a short time window. To control for uneven patterns of head rotation, the spike histogram was then divided by a summed, smoothed, histogram of all of the relative directional dwell times across all time windows: this defined the spatial cross-correlogram. To assess the degree of spatially coupled firing and the angle of directional offset, the spatial cross-correlogram (range −180° to 180° from the reference spike) was used to construct a polar rate map (analogous to that for normal head-directional firing), and the mean direction and mean resultant vector length (i.e. Rayleigh Vector) for this map were calculated. A large Rayleigh Vector indicates a strong spatial coupling over the time frame defined by the window. The degree of similarity between angular offsets in the standard and small boxes was compared using circular-circular regression. Only cell pairs for which both contributing cells were classified as HD cells in the small box, and neither were classified as HD cells in the standard box, were included in this analysis.

### Bayesian Decoding

Decoding in the standard box was performed using standard Bayesian methods (see for example [7, 8]) barring the key difference that the function of a neuron’s average firing with respect to direction (*f*_i_(**x**) in equation 35, ref [7]), was derived from firing rate maps recorded in the small box trial. The final Bayesian estimate of position:

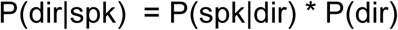

was therefore an estimate of the direction which current HD network activity was signalling, defined within the directional reference frame of the small box trial. See figure S1 for a comparison of this approach with the standard method, decoding standard box position using *f*_i_(**x**) based on standard box rate maps. Only ensembles containing at least 9 HD cells in at least one small box trial were included (N=6 ensembles). The decoding window was 1 sec long, and non-overlapping windows were used. To estimate the angular velocity at which the decoded direction was moving, the decoded position was up-sampled to 50Hz, smoothed with a 5 sec Gaussian kernel, and the angular velocity for each sample was estimated as the difference in direction at sample N and sample N+50 (i.e. across a 1 sec time step). To produce a comparable estimate of actual head angular velocity (in order to correlate this with the decoded estimate), the actual head direction of the animal was averaged across 1 sec long non-overlapping bins (mimicking the output of the decoding algorithm), then up-sampled and smoothed equivalently. The angular velocity was then estimated across 1 sec time steps. For small box decoding *f*_i_(**x**) was based on rate maps from the small box trial. In this case, odd minutes of the trial were used to decode even minutes of observed spiking, and vice versa, such that at no point was spiking data used to decode itself. The relationship between actual and decoded angular velocity was tested using linear regression, and the difference between the slopes of these regression fits (between small and standard box) was tested using student’s t, analogously to testing for the differences between two means (9). The rate of drift was defined as the change in the absolute offset between the actual and decoded head direction, calculated across 1 sec time steps, using decoded and actual head direction data that was up-sampled and smoothed as described for the estimation of angular velocity. Animals were defined as being close to walls if they were within 7.5cm of the closest wall, and were defined as being close to corners if they were less than 7.5cm radial distance from the closest corner.

### Estimation of coupled cell pairs in P12 data

To investigate the presence of temporally or spatially coupled neurons in P12 ensembles, we measured the proportions of temporal cross-correlograms with high or low central 1-sec mean normalised correlation (indicating temporally coupled or anti-coupled firing, respectively) and spatial cross-correlograms with high RV scores (indicating consistent spatial firing offsets with the timewindow). We defined 95% confidence limits for the chances of finding high or low coupling scores in non-HD cell data, by measuring the 5^th^ and 95^th^ percentiles of the distributions of scores in the population of all known nonHD-nonHD cell pairs that were recorded (N Pair=3,634). This population was based on all pairs of recorded neurons that were not classified as HD cells, under experimental conditions when stable HD cells could be detected (small box at ages P13-14; standard box and small box at ages P15-P21. The overrepresentation of coupling scores beyond these limits in P12 data (significantly greater than 5%) was tested using a 1-sample Z-test for proportions.

## Bibliography

1. Taube, J. S., Muller, R. U., and Ranck Jr., J. B. (1990). Head-direction cells recorded from the postsubiculum in freely moving rats. I. Description and quantitative analysis. J.Neurosci. 10, 420–435.

2. Finkelstein, A., Derdikman, D., Rubin, A., Foerster, J. N., Las, L., and Ulanovsky, N. (2014). Three-dimensional head-direction coding in the bat brain. Nature 517, 159–164.

3. Taube, J. S. (2007). The head direction signal: origins and sensory-motor integration. Annu.Rev.Neurosci. 30, 181–207.

4. Wills, T. J., Cacucci, F., Burgess, N., and O’Keefe, J. (2010). Development of the hippocampal cognitive map in preweanling rats. Science (80-. ). 328, 1573–1576.

5. Langston, R. F., Ainge, J. A., Couey, J. J., Canto, C. B., Bjerknes, T. L., Witter, M. P., Moser, E. I., and Moser, M. B. (2010). Development of the spatial representation system in the rat. Science (80-. ). 328, 1576–1580.

6. Mittelstaedt, H., and Mittelstaedt, M.-L.. (1982). Homing by path integration. In Avian Navigation, F. Papi and H. G. Wallraff, eds. (New York: Springer), pp. 290–297.

7. Etienne, A. S., and Jeffery, K. J. (2004). Path integration in mammals. Hippocampus 14, 180–192.

8. Yoganarasimha, D., Yu, X., and Knierim, J. J. (2006). Head direction cell representations maintain internal coherence during conflicting proximal and distal cue rotations: comparison with hippocampal place cells. J.Neurosci. 26, 622–631.

9. Taube, J. S., Muller, R. U., and Ranck Jr., J. B. (1990). Head-direction cells recorded from the postsubiculum in freely moving rats. II. Effects of environmental manipulations. J.Neurosci. 10, 436–447.

10. Skaggs, W. E., Knierim, J. J., Kudrimoti, H., and McNaughton, B. L. (1995). A model of the neural basis of the rat’s sense of direction. In Neural Information Processing Systems 7, S. J. Hanson, J. D. Cowan, and C. L. Giles, eds. (MIT Press), pp. 173–180.

11. Zhang, K. (1996). Representation of spatial orientation by the intrinsic dynamics of the head-direction cell ensemble: a theory. J.Neurosci. 16, 2112–2126.

12. Redish, A. D., Elga, A. N., and Touretzky, D. S. (1996). A coupled attractor model of the rodent head direction system. Network 7, 671.

13. Kim, S. S., Rouault, H., Druckmann, S., and Jayaraman, V. (2017). Ring attractor dynamics in the Drosophila central brain. Science 356, 849–853.

14. Green, J., Adachi, A., Shah, K. K., Hirokawa, J. D., Magani, P. S., and Maimon, G. (2017). A neural circuit architecture for angular integration in Drosophila. Nature 546, 101–106.

15. Peyrache, A., Lacroix, M. M., Petersen, P. C., and Buzsáki, G. (2015). Internally organized mechanisms of the head direction sense. Nat. Neurosci. 18, 569–75.

16. Hahnloser, R. H. (2003). Emergence of neural integration in the head-direction system by visual supervision. Neuroscience 120, 877–891.

17. Stringer, S. M., Trappenberg, T. P., Rolls, E. T., and de Araujo, I. E. T. (2002). Self-organizing continuous attractor networks and path integration: one-dimensional models of head direction cells. Network 13, 217–42.

18. Tan, H. M. H. M., Bassett, J. P. J. P., O’Keefe, J., Cacucci, F., and Wills, T. J. T. J. (2015). The Development of the Head Direction System before Eye Opening in the Rat. Curr. Biol. 25, 479–83. A

19. Bjerknes, T. L., Langston, R. F., Kruge, I. U., Moser, E. I., and Moser, M. B. (2015). Coherence among head direction cells before eye opening in rat pups. Curr. Biol. 25, 103–108.

20. Muessig, L., Hauser, J., Wills, T. J. T. J., and Cacucci, F. (2015). A Developmental Switch in Place Cell Accuracy Coincides with Grid Cell Maturation. Neuron 86, 1167–1173.

21. Hardcastle, K., Ganguli, S., and Giocomo, L. M. (2015). Environmental boundaries as an error correction mechanism for grid cells. Neuron 86, 827–39. Available at: http://www.ncbi.nlm.nih.gov/pubmed/25892299

22. Bassett, J. P., and Taube, J. S. (2001). Neural correlates for angular head velocity in the rat dorsal tegmental nucleus. J.Neurosci. 21, 5740–5751.

23. Sharp, P. E., Tinkelman, A., and Cho, J. (2001). Angular velocity and head direction signals recorded from the dorsal tegmental nucleus of gudden in the rat: implications for path integration in the head direction cell circuit. Behav.Neurosci. 115, 571–588.

24. Turner-Evans, D., Wegener, S., Rouault, H., Franconville, R., Wolff, T., Seelig, J. D., Druckmann, S., and Jayaraman, V. (2017). Angular velocity integration in a fly heading circuit. Elife 6.

25. Frohardt, R. J., Bassett, J. P., and Taube, J. S. (2006). Path integration and lesions within the head direction cell circuit: Comparison between the roles of the anterodorsal thalamus and dorsal tegmental nucleus. Behav. Neurosci. 120, 135–149.

26. Butler, W. N., Smith, K. S., van der Meer, M. A. A., and Taube, J. S. (2017). Erratum: The Head-Direction Signal Plays a Functional Role as a Neural Compass during Navigation (Current Biology (2017) 27(9) (1259–1267)(S0960982217303317)(10.1016/j.cub.2017.03.033)). Curr. Biol. 27, 2406.

27. Lannou, J., Precht, W., and Cazin, L. (1979). The postnatal development of functional properties of central vestibular neurons in the rat. Brain Res. 175, 219–232.

28. Stratton, P., Wyeth, G., and Wiles, J. (2010). Calibration of the head direction network: a role for symmetric angular head velocity cells. J. Comput. Neurosci. 28, 527–538.

29. McNaughton, B. L., Battaglia, F. P., Jensen, O., Moser, E. I., and Moser, M. B. (2006). Path integration and the neural basis of the “cognitive map.” Nat.Rev.Neurosci. 7, 663–678.

30. Fuhs, M. C., and Touretzky, D. S. (2006). A spin glass model of path integration in rat medial entorhinal cortex. J.Neurosci. 26, 4266–4276.

31. Burak, Y., and Fiete, I. R. (2009). Accurate path integration in continuous attractor network models of grid cells. PLoS.Comput.Biol. 5, e1000291.

32. Widloski, J., and Fiete, I. R. (2014). A Model of Grid Cell Development through Spatial Exploration and Spike Time-Dependent Plasticity. Neuron 83, 481–495.

33. Wills, T. J., Barry, C., and Cacucci, F. (2012). The abrupt development of adult-like grid cell firing in the medial entorhinal cortex. Front. Neural Circuits.

34. Muir, G. M., Brown, J. E., Carey, J. P., Hirvonen, T. P., la Santina, C. C., Minor, L. B., and Taube, J. S. (2009). Disruption of the head direction cell signal after occlusion of the semicircular canals in the freely moving chinchilla. J.Neurosci. 29, 14521–14533.

35. Hermer, L., and Spelke, E. S. (1994). A geometric process for spatial reorientation in young children. Nature 370, 57–59.

36. Cheng, K., Huttenlocher, J., and Newcombe, N. S. (2013). 25 years of research on the use of geometry in spatial reorientation: a current theoretical perspective. Psychon. Bull. Rev. 20, 1033–1054.

## References

1. Harris, K. D., Henze, D. A., Csicsvari, J., Hirase, H. & Buzsaki, G. Accuracy of tetrode spike separation as determined by simultaneous intracellular and extracellular measurements. J.Neurophysiol. 84, 401–414 (2000).

2. Wills, T. J., Cacucci, F., Burgess, N. & O’Keefe, J. Development of the hippocampal cognitive map in preweanling rats. Science (80-. ). 328, 1573–1576 (2010).

3. Langston, R. F. et al. Development of the spatial representation system in the rat. Science (80-. ). 328, 1576–1580 (2010).

4. Tan, H. M., Bassett, J. P., O’Keefe, J., Cacucci, F. Wills, T. J. The development of the head direction system before eye opening in the rat. Curr. Biol. 25, (2015).

5. Bonnevie, T. et al. Grid cells require excitatory drive from the hippocampus. Nat.Neurosci. 16, 309–317 (2013).

6. Chen, G., Manson, D., Cacucci, F. & Wills, T. J. Absence of Visual Input Results in the Disruption of Grid Cell Firing in the Mouse. Curr. Biol. 26, 2335–42 (2016).

7. Zhang, K., Ginzburg, I., McNaughton, B. L. & Sejnowski, T. J. Interpreting neuronal population activity by reconstruction: unified framework with application to hippocampal place cells. J.Neurophysiol. 79, 1017–1044 (1998).

8. Muessig, L., Hauser, J., Wills, T. J. T. J. & Cacucci, F. A Developmental Switch in Place Cell Accuracy Coincides with Grid Cell Maturation. Neuron 86, 1167–1173 (2015).

9. Zar, J. H. Biostatistical Analysis. (Prentice Hall, 2010).

